# Pot1 promotes telomere DNA replication via the Stn1-Ten1 complex in fission yeast

**DOI:** 10.1101/2023.01.23.525167

**Authors:** Pâmela C. Carvalho Borges, Jose Miguel Escandell, Samah Matmati, Stéphane Coulon, Miguel Godinho Ferreira

## Abstract

Telomeres are nucleoprotein complexes that protect the chromosome-ends from eliciting DNA repair while ensuring their complete duplication. Pot1 is a subunit of telomere capping complex that binds to the G-rich overhang and inhibits the activation of DNA damage checkpoints. In this study, we explore new functions of fission yeast Pot1 by using a *pot1-1* temperature sensitive mutant. We show that *pot1* inactivation impairs telomere DNA replication resulting in the accumulation of ssDNA leading to the complete loss of telomeric DNA. Recruitment of Stn1 to telomeres, an auxiliary factor of DNA lagging strand synthesis, is reduced in *pot1-1* mutants and overexpression of Stn1 rescues loss of telomeres and cell viability at restrictive temperature. We propose that Pot1 plays a crucial function in telomere DNA replication by recruiting Stn1-Ten1 and Polα-primase complex to telomeres, thus promoting lagging-strand DNA synthesis at stalled replication forks.

## Introduction

Telomeres, the ends of eukaryotic chromosomes, contain short double-stranded tandem repeats that terminates in a single-stranded DNA (ssDNA) overhang. They are bound by a protein complex, termed Shelterin, which ensures their protection (1–3). In humans, Shelterin is composed by TRF1 and TRF2 that directly bind to the telomeric duplex DNA and recruit RAP1. Telomeric ssDNA is bound by POT1, that recruits TPP1, and the two subcomplexes are bridged by TIN2. Shelterin prevents inappropriate DNA repair at chromosome-ends, checkpoint activation and facilitates replication of telomeric DNA (4, 5). Telomeres shortening occurs with successive cell divisions caused by the inability of replicative DNA polymerases to fully replicate terminal DNA. This process is known as “end replication problem” (6). The slow telomere erosion is counteracted by the action of telomerase, a reverse transcriptase that uses the telomere 3’ overhang to extend chromosome-ends (1, 7). However, abrupt loss of telomeric DNA can occur during telomeric DNA replication. Indeed, telomeres are genome fragile sites hard to replicate as they contain several DNA replication barriers (DNA secondary structures, R-loop containing RNA:DNA hybrids, bound proteins and topological constrains). These barriers impede the progression of the replication fork through telomeric repeat sequences leading to fork collapse and, potentially, DNA double strand breaks (DSBs) (8, 9). Thus, unidirectional replication forks that progress toward the end of the chromosome frequently slow down and stall when entering telomeric tracks (10, 11).

The fission yeast *Schizosaccharomyces pombe* contains a shelterin-like complex that is evolutionary conserved between fission yeast and humans (12). Taz1, the homolog of both mammalian TRF1 and TRF2, binds to duplex telomeric DNA wherease the 3’ overhang is protected by Pot1. Both factors are physically bridged by Rap1, Poz1 and Tpz1 and ensure the full protection of telomeres (13, 14). *S. pombe* shelterin plays a major function in recruiting telomerase that includes the catalytic subunit Trt1 and the RNA subunit *ter1*. Phosphorylation of Shelterin by the Rad3^ATR^ kinase promotes the recruitment of telomerase to fission yeast telomeres via Tpz1 (17, 18) resulting in telomere elongation (19). The cycle of elongation is terminated through the sumoylation of Tpz1, which promotes the recruitment of the Stn1-Ten1 complex (20–22). Stn1-Ten1 fulfils an important function in telomere protection (25) by recruiting Polα-Primase complex and lagging strand synthesis (22, 23).

As in mammals, fission yeast Shelterin regulates telomere replication. Taz1 is a major factor in this process although its precise function has not yet been established (10). Additional components, such as the RPA heterotrimer, the Pfh1 helicase and telomerase itself, contribute to promoting efficient replication of telomeres (24–27). Recently, we and others have shown that the Stn1-Ten1 complex is required for telomere and subtelomere replication (22, 28). Stn1-Ten1 prevents the accumulation of telomeric ssDNA by promoting lagging strand DNA synthesis through thecruitment of Polα-Primase complex to stalled replication forks. In mammals, the CST complex is composed by CTC1, STN1 and TEN1 and is not only required for telomere replication (29–31) but also global genome replication (32, 33). The CST complex may also act as a terminator of telomerase elongation by promoting lagging strand synthesis and blocking the 3’-overhang (34, 35).

Pot1 is an essential subunit of the Shelterin complex. It recognizes the 3’ single-stranded overhang and plays a major role in the protection of telomeres (36, 37). This function is reinforced by its interaction with Tpz1 (38). Pot1 is also described to act as a negative regulator of telomerase (39). Indeed, mammalian POT1 is thought to promote telomere lagging strand synthesis by the recruitment of the CST complex to telomeres (40, 41). Thus, functions of POT1 seems to be intimately linked to CST complex. However, in contrast to CST, a role of POT1 in telomere replication has not been established so far. A thermo-sensitive (*ts*) mutant of *pot1*^*+*^ (*pot1-1)* was identified by the Cooper laboratory (42). This mutant rapidly looses telomeres and accumulate ssDNA during S-phase at restrictive temperature, indicating that Pot1 fulfils crucial functions during DNA replication (42). Using *pot1-1* mutants, we further explore the role of Pot1 in telomere replication. We show that Pot1 acts with the Stn1-Ten1 complex to promote DNA replication of telomeric and subtelomeric DNA. Our results indicate that Pot1 stabilizes Stn1-Ten1 at telomeres and promotes Polα−dependent DNA synthesis to ensure efficient replication of chromosome ends.

## Materials and Methods

### Yeast strains and media

The strains used during this work can be found listed in supplementary table1. *De novo* strains were constructed using commonly used techniques (43). These include yeast mating combined with tetrad dissection and yeast transformation. Intended mutants were screened using appropriate selective media combined with Polymerase Chain Reaction (PCR) tests. Gene knockouts were made by substituting entire Open Reading Frames (ORF) with kanMX6, hphMX6 or natMX6 cassettes while gene tagging was made using GFP, HA or Myc epitopes. *pot1-1* strain generated in the Cooper lab resulted from integration of GFP-KanMX cassette at the C-terminal part of endogenous *pot1+* carrying E158G and D456N mutations (42).

Standard media and growth condition were used, unless stated otherwise. The growth media used for most of the strains was Yeast Extract Supplemented (YES) and the normal growth temperature was 32°C. Temperature sensitive strains were cultured at 25°C for non-conditioned protein (permissive temperature) and at 37°C to inactivate the protein being studied (restrictive temperature). Strains that have genes under *nmt* promoter were grown in Pombe Minimal Glutamate medium (PMG), which lacks thiamine to induce overexpression of the target gene. The media was complemented with the required amino acids.

### Spot viability assay

Cells were grown in YES or PMG liquid media in a shacking incubator (200 rpm) until mid-logarithmic growth phase is reached (OD≈ 0.7). Incubation temperature was 32°C while temperature sensitive strains were incubated at 25°C. 5-fold serial dilution were made from 0.7 × 10^6^ cells and each dilution were spotted in YES or PMG solid media. Plates were incubated at 25°C to control for the loading while others were incubated at 37°C for 3-5 days before imaging.

### Microscopy and Cell length Measurement

Cells were grown in liquid media at 25°C for temperature sensitive strains (permissive temperature) and 32°C for remaining strains until mid-logarithmical growth phase. They were then incubated in a 37°C shaking water bath. Eight hours after temperature shift cells were fixed in 70% ethanol and imaged using Delta Vision Core System Microscope (100x 1.4 numerical aperture UplanSApo objective and a cascade2 EMCCD camera). Deconvolution was performed using the enhanced ratio method in softWoRx software. Lengths of 400 cells per strain were assessed using FIJI Software and analyzed using PRISM Software.

### Southern Blotting

Cells were grown in liquid media and genomic DNA was extracted from exponentially growing yeast cells using phenol-chlorophorm technique. DNA samples were digested using *EcoR*I enzyme and resolved by electrophoresis in a 0.7% agarose gel, which were treated for 15 minutes in 0.25M HCl, 30 minutes in denaturing solution (0.5M NaOH, 1.5M NaCl) and 1-hour in the neutralizing solution (1M NH4Ac, 20mM NaOH). DNA was transferred from the agarose gel to a positively charged membrane (Hybond-XL membranes from GE healthcare) using capillarity technique. After being exposed to UV radiation to induce DNA cross-link the membrane was pre-hybridized in Church-Gilbert solution (1% BSA, 1 mM EDTA, 7% SDS, 0.5 M Na2HPO4, 4 ml H3PO4 85%, up 1litter with H2O) at 65°C and probed using Telomere oligo radiolabeled with radioactive phosphate ^32^P to visualize the telomeres. To measure the loading the membrane was probe with radiolabeled Rad4 oligo that was created using 5’-tcataatttcccccaaacca-3’ and 5’ accgtattttagccgacgtg-3’ primers. We also probed subtelomeric region that is 5Kbp from telomere using ^32^P-labeled PCR products amplified with 5΄-acactcaattcaaatcaacttc-3΄ and 5΄-gtgtttgaaaattgagcttatg-3΄ primers.

### Overhang Assay

Yeast cultured in liquid media was used to extract DNA using phenol-chlorophorm technique after Zymolyase treatment. In-gel telomeric overhang assay was performed on *EcoR*I-HF digested DNA samples. DNA was resolved by electrophoresis in a 0.7% agarose gel which was dried for two hours at 50°C using 583 gel dryer model from Bio-Rad. Dried gel was hybridized in Church-Gilbert solution (1% BSA, 1 mM EDTA, 7% SDS, 0.5 M Na2HPO4, 4 ml H3PO4 85%, up 1litter with H2O) at 50°C for one hour and probed overnight using telomeric C-rich oligo radiolabelled with γ-ATP.

### Two-Dimensional (2D) gel electrophoresis

2D gel electrophoresis assay was implemented as described by Noguchi *et al* (44). 60 U of *Nsi*I enzyme was used to digest 10 μg of DNA which was used to access telomeres. The first dimension DNA was run on 0.4% agarose gel while the second dimension was run in a 1% agarose gel. Gels were transferred to positively charged membranes (Hybond-XL membranes) and probed with the STE1 probe.

### Pulse Field Gel Electrophoresis (PFGE)

Cells cultured in liquid media were collected when they reached mid-logarithmic phase, washed twice with SP1 (1.2 M d-sorbitol, 50 mM sodium citrate, 50 mM, Na2HPO4·7H2O, 40 mM EDTA pH6.5) and treated for 40 minutes with Zymolase-20T (30mg/100 μl). Spheroplasts were then spun down and re-suspended in 1% low melting point agarose in TSE (10mM Tris-HCl pH7.5, 0.9M sorbitol, 45mM EDTA) to have a final concentration of 1×10^8^ cells in 100ul. The mix was dispensed in 90 μl plugs molds and let to solidify at 4°C. Plugs were then incubated for 90 minutes in 0.25M EDTA, 50mM Tris-HCl pH 7.5 and 1% SDS and then in 0.5M EDTA 10mM Tris (pH9.5), 1% lauryl sarcosine and 1-2mg/ml proteinase K for 24 hours at 55°C. Three washes of 30 minutes in 10mM EDTA and 10mM Tris-HCl at 25°C were made and then plugs were treated for one hour with 10mM EDTA,10mM Tris-HCl and 0.04mg/ml PMSF at 55°C. Three other 30 minutes washes were made with 10mM EDTA and 10mM Tris-HCl at 25°C and plugs were digested over night with *Not*I enzyme after being treated with *Not*I buffer for one hour at 37°C. We loaded the plugs into 1% pulse-field certified agarose gel dissolved in 0.5X TBE and used CHEF Mapper PFGE System to run the gel at 14°C, 6V/cm with pulse time of 60 to 120s. We then transferred the DNA to a positively charged membrane (Hybond-XL membranes), pre-hybridized with Church-Gilbert solution (1% BSA, 1 mM EDTA, 7% SDS, 0.5 M Na2HPO4, 4 ml H3PO4 85%, up 1L with H2O) at 65°C for one hour and probed using Telo and LMAC oligos radiolabeled with ^32^P.

### Chromatin Immunoprecipitation (ChIP)

Protocol described by Moser *et al*. (45) was used to perform ChIP assay. Cells at mid-log growth phase were collected and fixed with 11% formaldehyde (dilution was made with 0.1M NaCl, 1mM EDTA, 50mM HEPES-KOH, pH 7.5) for 20 min at room temperature. 0,25M glycine was used to quench the solution for 5 minutes. Cells were pelleted, washed twice with cold PBS and lysed with 2x lysis buffer (100 mM Hepes-KOH, pH 7.5 2 mM EDTA 2% Triton X-100 0.2% Na Deoxycholate) by mechanical disruption method. QSonica Sonicator Q800R3 was used to shear the chromatin and then equal amounts of DNA were used for immunoprecipitation protocol with anti-Myc (9E10; Santa Cruz biotechnology) with magnetic Protein A beads. The DNA-protein complexes were washed with lysis buffer 0.1% SDS/275 mM NaCl, lysis buffer 0.1% SDS/500 mM NaCl and last with 10 mM Tris-HCl, pH 8.0, 0.25M LiCl, 1 mM EDTA, 0.5% NP-40, 0.5% Na Deoxycholate solution. DNA was precipitated using sodium acetate, 2-Propanol and glycogen. DNA was resuspended with 50 mM Tris-HCl, pH 7.5, 10mM EDTA, 1% SDS, de-crosslinked, purified used to in triplicate together with SYBR-Green-based to perform a quantitative real-time PCR (Bio-Rad) using the primers described in Carneiro, *et al*. (46).

## Results

### Pot1 inactivation prevents Stn1 recruitment to telomere

Deletion of *S. pombe pot1*^*+*^ leads to rapid telomere loss and survival through chromosome circularization (32). To circumvent this terminal phenotype, we used a *pot1* temperature sensitive mutant previously described by the group of J. Cooper (*pot1-1*) (42). At the permissive temperature (25°C), *pot1-1* cells grow normally although with slightly longer telomeres (42). When temperature is raised to 37°C (restrictive temperature), *pot1* function is inactivated and viability of *pot1-1* mutant is severely reduced (**Figure 1A**). In fission yeast, checkpoints delay cell cycle progression resulting in cell elongation (47). To investigate if *pot1-1* reduced viability at 37°C was due to checkpoint activation, we measured cell length and observed that, at permissive temperature, WT and *pot1-1* cells length averaged 10μm (**Figure 1B**). However, when the temperature was raised to 37°C for 8 hours, *pot1-1* elongated to an average length of 20μm (**Figure 1B**). As previously shown (42), *pot1-1* cells arrest at the restrictive temperature through a DNA-damage response (DDR) pathway. To assess the telomeric effects of *pot1-1* inactivation, we performed a Southern blot probing for telomeric sequences. Whereas *pot1-1* have longer telomeres than WT at 25°C, after 8 hours of incubation at 37°C, we observed a significant loss of telomeres sequences, revealed by the disappearance of telomeric signal (**Figure 1C**), confirming previous results by the Cooper lab (42). To investigate if *pot1-1* inactivation led to chromosomes circularization as *pot1Δ*, we performed a Pulse Field Gel Electrophoresis (PFGE) of *NotI*-digested genomic DNA followed by Southern blotting (**Figure 1D**). In contrast to telomerase mutants (*trt1Δ*) and to *pot1Δ* mutant that circularize their chromosomes (48, 49), we were unable to detect chromosome fusion bands (L+I and C+M) in *pot1-1* mutants, indicating that chromosomes remained linear after 8 hours at 37°C despite the inability to detect telomere sequences.

**Figure 1.**
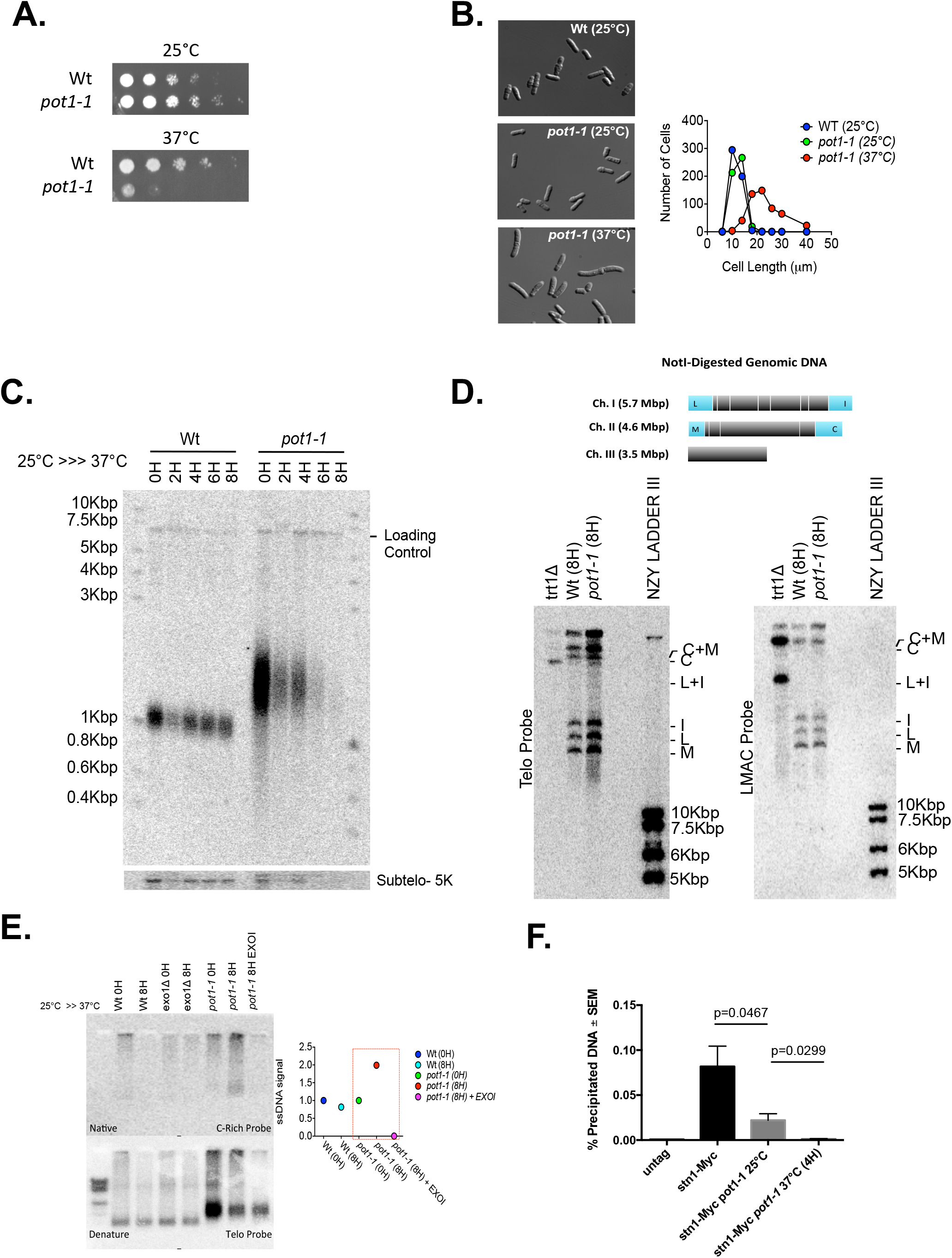
Pot1 inactivation reduces Stn1 recruitment to telomeres. **A.)** *pot1-1* inactivation compromises cell viability. Cell viability was assessed through spot dilution assay. **B.)** Cell elongation is triggered upon *pot1-1* inactivation. Delta Vision Core System were used to image the cells and their length were assessed using FIJI software. **C.)** *pot1-1* deficiency leads to rapid telomere loss. Chromosomes loose telomeres upon *pot1-1* inactivation. Cells were collected at 0H, 2H, 4H and 8H after temperature shift and DNA samples were digested with *EcoR*I enzyme. **D.)** Pulse Field Gel Electrophoresis shows no chromosome circularization upon eight hours of *pot1-1* inactivation. Samples were digested with NotI enzyme and Southern blots were radiolabeled with telomeric and LMIC probes. Mutants for telomerase (*trt1*∆) controlled for cells containing circular chromosomes. **E.)** *pot1-1* cells have longer G-rich overhang when Pot1 is inactive. In-gel hybridization assay was performed to access telomeric ssDNA. Samples were digested with *EcoR*I enzyme and radiolabeled with C-rich probe when native and telomeric probe when denatured. Mutants for telomerase (*exo1*∆) controlled for cells deficient for telomere end-resection. Exogenous exonuclease (EXOI) enzyme was able to digest all the ssDNA of *pot1-1* samples collected at the restricted temperature. **F.)** Stn1 recruitment to *pot1-1* telomere is significantly reduced at permissive temperature and Pot1 inactivation abolishes Stn1-myc binding to telomeres. ChIP analysis was used to quantify Stn1-myc recruitment to telomere. n ≥ 3; \**p* ≤0.05 based on a two-tailed Student’s t-test to control sample. Error bars represent standard error of the mean (SEM).

In fission yeast, the presence of RPA-coated ssDNA induces Rad3-dependent checkpoint activation (50). Because *pot1-1* cells activate DDR at restrictive temperature, we hypothesized that *pot1* deficiency leads to an increase in telomeric ssDNA. Thus, we performed native in-gel hybridization assay with a C-rich telomeric probe to detect the G-rich strand. We observed that *pot1-1* inactivation led to an increased in telomeric ssDNA (**Figure 1E**). We treated *pot1-1* genomic DNA with EXOI, an exogenous exonuclease that degrades linear ssDNA in the 3’ to 5’ direction. As seen on native gels, telomeric ssDNA of *pot1-1* grown at 37°C was lost upon treatment with EXOI (**Figure 1E**). These observations indicate that telomeric G-rich ssDNA formed upon *pot1* inactivation exhibits a 3’end.

The Stn1-Ten1 complex was shown to inhibit telomerase binding at telomeres and to promote lagging-strand synthesis through its interaction with Tpz1 and the recruitment of Polα-Primase (20–23). Thus, we analyzed the pattern of Stn1 recruitment to telomeres in *pot1-1* cells by performing a Chromatin Immunoprecipitation (ChIP). ChIP of myc-tagged Stn1 revealed a significant reduction of Stn1 present at telomeres of *pot1-1* already at the permissive temperature (**Figure 1F**). This reduction was further exacerbated by growing *pot1-1* cells for 4 hours at the restrictive temperature (**Figure 1F**). The observation that Stn1 recruitment to telomeres was significantly reduced in *pot1-1* mutants, even at the permissive temperature, shows that Pot1 is important for Stn1 recruitment to telomeres. At restrictive temperature, the absence of Stn1 binding at telomeres is likely to be a compound result from both Pot1 inactivation and loss of telomeric DNA. Importantly, the reduction of Stn1 binding to telomeres at the permissive temperature can explain the presence of longer telomeres in *pot1-1*. Reduction of Stn1-Ten1 complex results in the retainment of telomerase at telomeres and longer telomere extension. The same phenotype can observed in *stn1* mutants (see below). Although no direct interaction has been detected between Pot1 and Stn1 (or Ten1) (51), our data suggests that Pot1 participates to Stn1 recruitment to telomeres, likely by stabilizing Tpz1 and favoring Tpz1-Stn1 interaction.

### pot1 and stn1 function in the same genetic pathway for telomere length maintenance

Like *pot1-1* mutants, *stn1-1* temperature sensitive mutants have longer telomeres than WT even at permissive temperature (25°C) and its inactivation results in loss of telomeres and cell cycle arrest at restrictive temperature (28). In addition, *stn1* inactivation leads to increased 3’overhang extension (28). Taking into consideration these striking similarities and the observation that Pot1 is important for Stn1 recruitment to telomeres, we hypothesized that Pot1 controls Stn1 function in telomere maintenance, possibly acting upstream of Stn1. To test this hypothesis, we generated a *pot1-1 stn1-1* double mutant. At 25**°**C, WT and mutants grew similarly well, while at restrictive temperature the combination of both alleles is synthetically lethal (**Figure 2A**). Confirming the synergistic effect on growth, *pot1-1 stn1-1* cells elongated up to 30μm while the single mutants ranged from 15 to 20μm when grown at 37**°**C for 8 hours (**Figure 2B**). The combination of *pot1-1* and *stn1-1* mutations on telomere length appeared to be additive as derived from crossing both parental strains. *pot1-1 stn1-1* cells displayed a compound of longer telomeres similar to *stn1-1* single mutants and shorter telomeres like *pot1-1* single mutants (**Figure 2C-D)**. This result indicates that *pot1* and *stn1* are epistatic regarding the regulation of telomere length. However, *pot1-1 stn1-1* cells lose their telomeres at restrictive temperature faster than either single mutant (**Figure 2C**). Thus, in contrast, *pot1* and *stn1* appear to participate independently in telomere protection, not being able to compensate for the loss of each other’s deficiency. Taken together, our data indicate that *pot1* and *stn1* function in the same pathway in telomere length regulation and that both fulfil independent functions to protect telomeric DNA.

**Figure 2.**
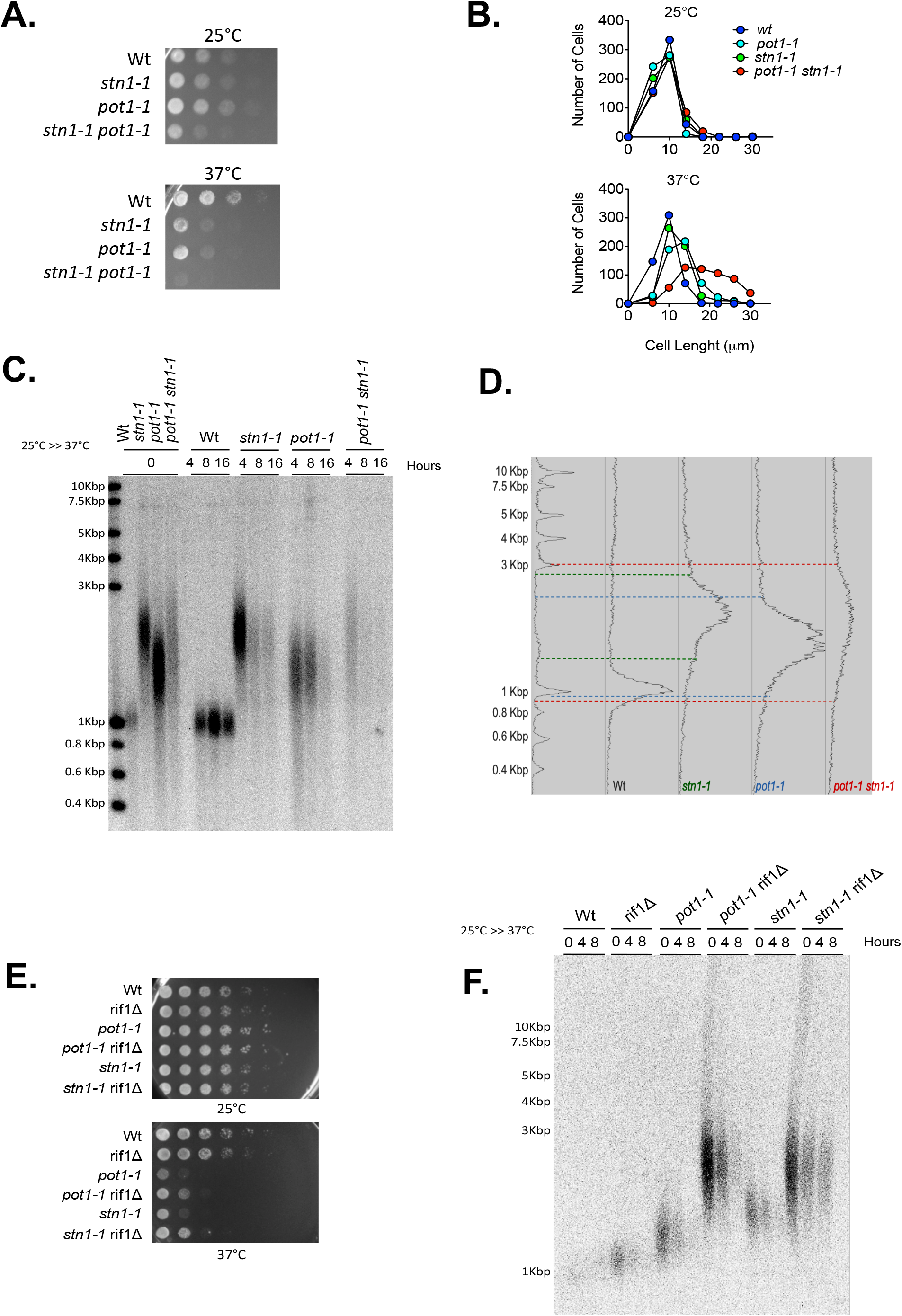
*pot1-1* and *stn1-1* are epistatic for telomere length but not for cell viability. **A.)** Spot Assay revealed that *pot1-1 stn1-1* double mutant cells display lower cell viability upon inactivation than either *pot1-1* or *stn1-1* single mutants. **B.)** *pot1-1 stn1-1* double mutants elongate more at the restrictive temperature than either *pot1-1* or *stn1-1* single mutants. **C.)** *pot1-1* and *stn1-1* are epistatic for telomere length and telomere loss. Southern blot was performed using samples collected at 0H, 4H, 8H and 16H after temperature shift to 37°C. DNA samples were digested with EcoRI enzyme and the membrane was radiolabeled with telomeric probe. **D.)** Spot viability assay revealed a rescue of Pot1 and Stn1 deficiency upon *rif1* deletion. **E.)** Southern blotting shows that *rif1* deletion rescues telomere loss that characterizes Pot1 and Stn1 inactivation. Samples were digested with *EcoR*I enzyme and membrane radiolabeled using a telomeric probe.

Previous studies showed that *rif1*^*+*^ deletion restored *stn1-1* viability and reduces the rate of telomere loss at the restrictive temperature (22, 28). Similarly, we wondered if the lack of Rif1 could also rescue *pot1-1* deficiency. Thus, and we combined *rif1*Δ deletion with *pot1-1* allele. Strikingly, we observed a rescue of *pot1-1* growth defects by *rif1*Δ mutants **(Figure 2E**). In addition, *rif1*Δ partially suppressed the loss of telomeres of *pot1-1* cells at restrictive temperature (**Figure 2F**). These observations suggest that the function of Rif1, known to repress subtelomeric replication origin firing, is deleterious for *pot1-1* cells, further linking Pot1 to telomere replication.

### Pot1 promotes efficient replication of telomeres

Because loss of telomeric DNA occurs in S-phase in *pot1-1* cells (42) and our observed genetic interactions between *rif1*Δ and *pot1-1*, we investigated the replication dynamics of *pot1-1* using two-dimensional gel electrophoresis (2D-gel) at telomeres, as previously described (10, 22, 23). Genomic DNA samples, from the parental WT and *pot1-1*, grown at permissive and restrictive temperatures, were analyzed after *Nsi*I digestion by Southern blotting using a subtelomeric (STE1) probe (**Figure 3A**). In the first dimension, we observed four distinct bands for the parental WT (*pot1+)* whereas *pot1-1* mutants possessed only one thick band (**Figure 3A**). This suggests that subtelomeric regions of *pot1-1* are highly rearranged, reminiscent to *taz1*Δ, *ss72*Δ and *stn1-226* mutants (10, 22, 23). Y-structures, corresponding to the unidirectional moving of the replication forks within the *Nsi*I fragment, were visualized in the WT strain at both temperatures (**Figure 3B**). However, these structures were not detected in *pot1-1* cells in either condition (**Figure 3B**). We concluded that replication of telomeric regions is severely affected in *pot1-1* at 25°C, leading to superimposed Y arcs that spread into higher molecular weight intermediates, mimicking the phenotype observed in *taz1*Δ (10). At restrictive temperature, the replication intermediates of *pot1-1* were barely detected, consistent with the disappearance of telomeric DNA in these conditions. Thus, from the 2D-gel analysis, we conclude that Pot1 is actively involved in the replication of telomeric DNA, possibly by facilitating Stn1 recruitment and its role in DNA end replication.

**Figure 3.**
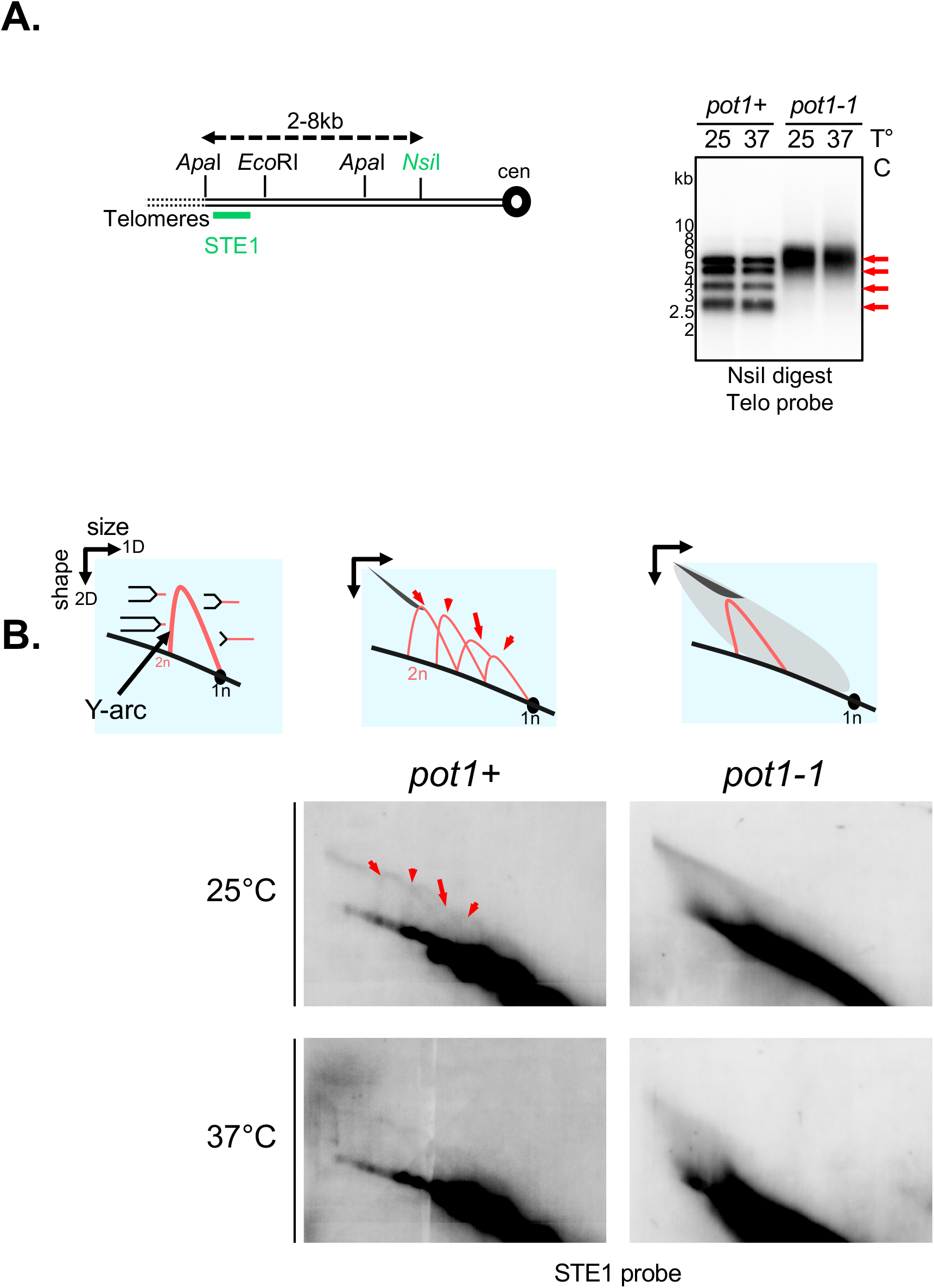
Pot1 inactivation induces replication fork stalling and collapse. **A.)** Relative position of the restriction sites in the subtelomeric regions of chromosomes I and II. The subtelomeric probe (STE1) that is used for 2D-gel hybridization (see below) is represented. Cen, centromere. Southern blot analysis of *Nsi*I telomeric fragments (first dimension) from the parental Wt strain and *pot1-1*mutant revealed by STE1 probe. **B.)** 2D-gel analysis of *Nsi*I telomeric fragments of Wt and *pot1-1* strains at 25°C and after 4 hours at 37°C. The Y-arc pattern is generated by unidirectional movement of a replication fork across each telomeric fragment shown in the first dimension. The cone-shaped signal represents four-way DNA junctions (double Y).

### Over-expression of *stn1*^*+*^ rescues *pot1-1* deficiency

Our data indicate that *pot1* and *stn1* are crucial for telomere replication and that Pot1 is important to recruit Stn1 to telomeres. Thus, we hypothesized if overexpression of *stn1*^*+*^ could restore defects of *pot1-1* mutants. To test this, we replaced the endogenous promoter of *stn1*^*+*^ by the *nmt* promoter (i.e., *nmt1-stn1*) and incorporated it in *pot1-1* mutants. Strikingly, overexpression of *stn1*^*+*^ restored the growth defect of *pot1-1* mutants at the restrictive temperature (**Figure 4A**). Moreover, this growth rescue at 37°C depended on the strength of the *nmt* promoter that controlled the level of expression of *stn1*^*+*^ (**Figure S1**). Indeed, expression of Stn1 with weaker versions of the *nmt* promoter (*nmt41* and *nmt81)* restored *pot1-1* viability only partially (**Figure 4A**). Consistent with this, overexpression of *stn1*^*+*^ also abrogated cell elongation of *pot1-1* at 37°C (**Figure 4B**). Next, we assessed the impact of the overexpression of *stn1*^*+*^ on telomere maintenance. We found that higher expression of *stn1*^*+*^ prevented both the loss of telomeric DNA (**Figure 4C**) and accumulation of telomeric ssDNA after 8 hours at 37°C (**Figure 4D**). Conclusively, *pot1-1* mutants longer telomeres were also rescued by *stn1*^*+*^ overexpression (**Figure 4C**). Taken together, our data indicates that Pot1 regulates telomere length and replication by the recruitment of Stn1 to telomeres.

**Figure 4.**
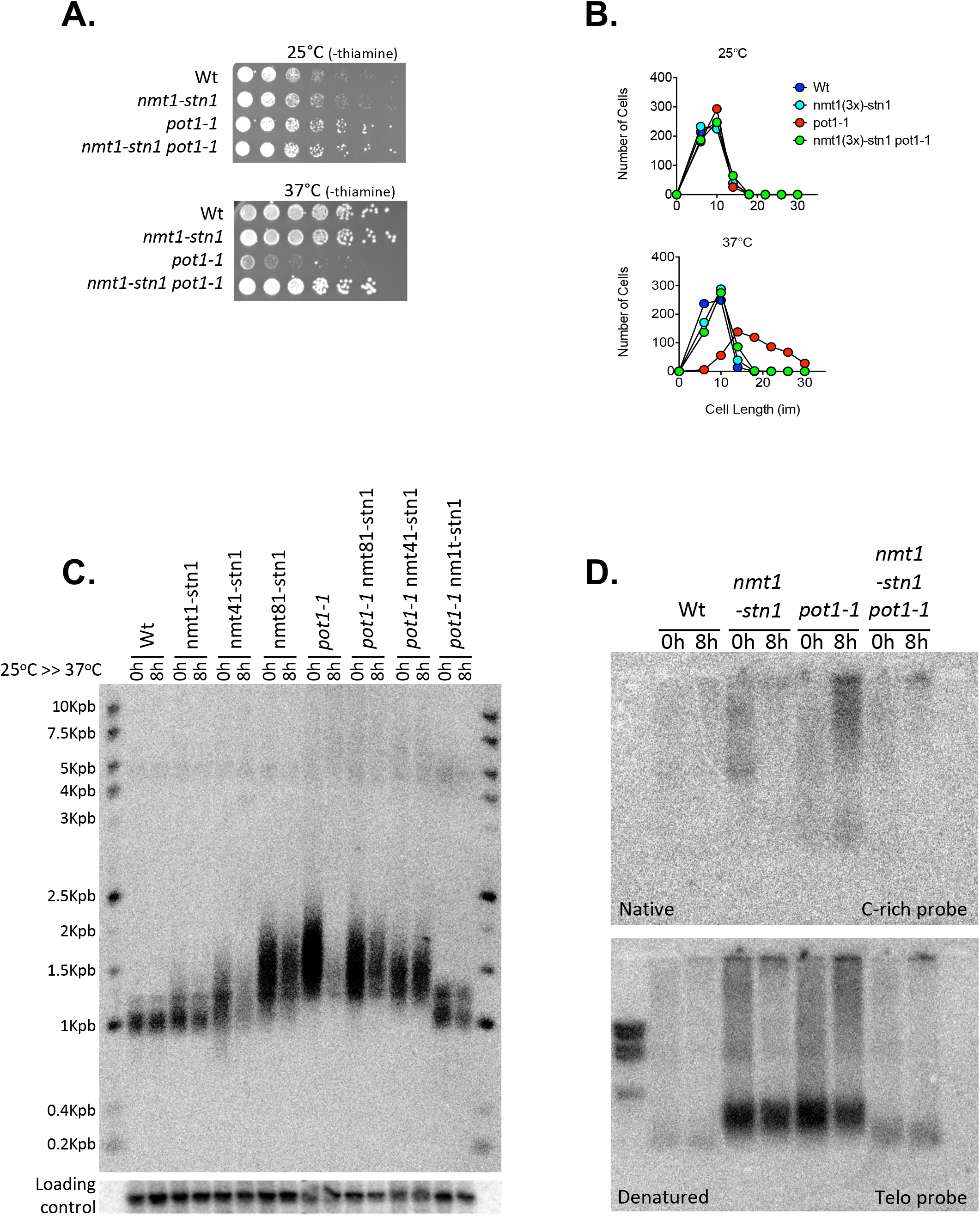
Overexpression of *stn1*^*+*^ rescues *pot1-1* deficiencies. **A.)** Spot viability assay shows that *stn1*^*+*^ overexpression rescues *pot1-1* cells viability. The *nmt1* promoter was used to induce *stn1*^*+*^ overexpression in both Wt and *pot1-1* cells. **B.)** Cell length measurements reveals that *stn1*^*+*^ overexpression rescues *pot1-1* cell lengthening at the restrictive temperature. **C.)** *stn1*^*+*^ overexpression rescues telomere length and telomere loss of *pot1-1* cells to Wt levels. To assess the length of telomeres we performed a Southern blotting with samples collected at 0H and 8H after *pot1-1* inactivation. **D.)** *stn1*^*+*^ overexpression prevents the accumulation of telomeric ssDNA upon Pot1 inactivation. In-gel hybridization using *EcoR*I digested sample were radiolabeled with C-rich probe for native gels, while telomeric probe was used for denatured gels.

### Over-expression of Polα partially rescues *pot1-1* deficiency

The (C)ST complex promotes DNA replication through the recruitment of the Polα-primase complex at telomeres and stalled replication forks (52). We previously showed that the overexpression of the catalytic subunit of the DNA polymerase α (Pol1) partially restores viability and telomere loss of *stn1* mutants (22). In line with this, we tested whether overexpression *pol1*^*+*^ driven by pREP41-Pol1 plasmids could also rescue the defects of *pot1-1* mutants. Indeed, we observed that overexpression of *pol1*^*+*^ partially restores the growth defects of *pot1-1* mutant at restrictive temperature (**Figure 5A**), and partially prevents the loss of telomeric sequences (**Figure 5B**), thus paralleling the positive effects observed on *stn1* mutants (22). Thus, our data suggests that Pot1 facilitates re-initiation of DNA synthesis at stalled or collapsed replication forks through recruitment and/or the stabilization of Stn1-Ten1 complex at telomeres leading to fork restart by the Polα-primase complex.

**Figure 5.**
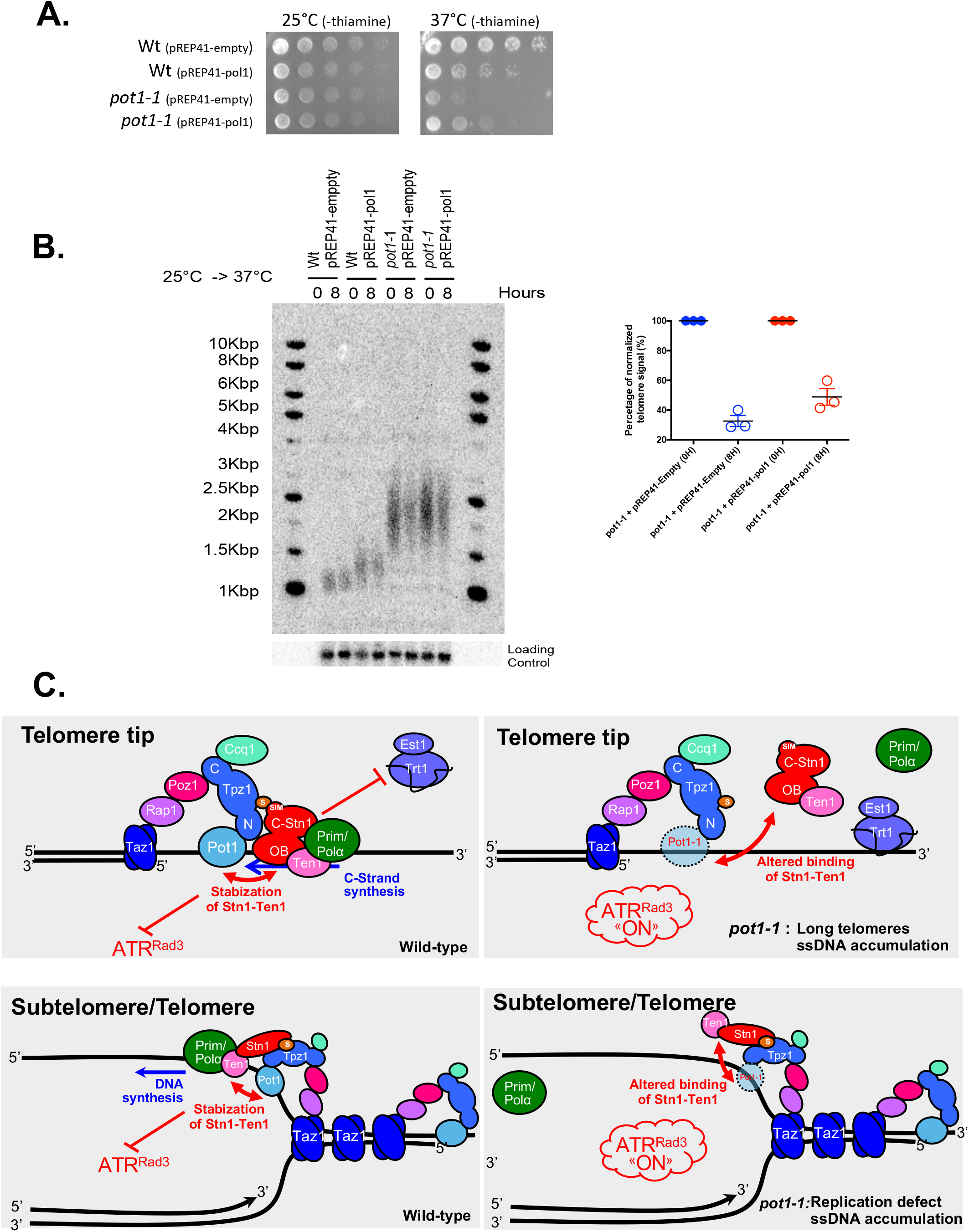
Polα overexpression rescues cell viability and telomere loss of *pot1-1* cells. **A.)** Spot viability assay reveals that *pol1*^*+*^ overexpression partially rescues the viability of *pot1-1* cells at 37°C. **B.)** Quantification of Southern blot shows a modest rescue of *pot1-1* telomere loss 8H after Pot1 inactivation. DNA samples were digested with *EcoR*I enzyme and membrane was radiolabeled with telomeric probe. **C.)** Dual function of Pot1 at telomeres *via* the recruitment of Stn1-Ten1. We showed here that Pot1 is crucial to recruit and/or stabilize Stn1-Ten complex at telomeres. Through this function, we propose that Pot1 has a dual function at telomeres, first in regulating telomere length and, second, by participating in telomere DNA replication. Both functions rely on the ability of Pot1 to recruit Stn1-Ten1 complex and to promote DNA synthesis by Polα-primase, explaining how Pot1 limits the accumulation of ssDNA at telomeres and prevents the activation of ATR-dependent DDR. When Pot1 is impaired, Stn1-Ten1 recruitment is reduced and telomerase inhibition is altered leading to the lengthening of telomeres. This also causes the accumulation of ssDNA and DDR activation through ATR^Rad3^. Telomere DNA replication is also hindered by Pot1 inactivation. This also leads to ssDNA accumulation during DNA replication and DDR activation.

## Discussion

Pot1 is an essential subunit of the Shelterin telomere binding complex that binds directly the telomeric single-stranded DNA and to Tpz1 (2). A main function of Pot1 is to protect chromosomal ends from an inappropriate DNA damage response, by blocking the accumulation of RPA-coated ssDNA, thereby inhibiting the recruitment of Rad3^ATR^ (37, 53). The (C)ST complex interacts with Tpz1/Pot1 that both promote lagging strand DNA replication (41) and negatively regulate telomerase (39). In this study, we further explored the role of Pot1 in fission yeast using *pot1-1* mutant. Our data demonstrate that Pot1 has a dual role at chromosomes ends. First, in regulating telomere length and, second, by participating in telomere DNA replication (**Figure 5C**). Both functions rely on the ability of Pot1 to recruit Stn1-Ten1 complex and to promote DNA synthesis by Polα-primase. This mechanistically explains how Pot1 limits the accumulation of ssDNA at telomeres and prevents Rad3^ATR^-dependent DDR. Although Pot1 does not appear to bind directly to Stn1 or Ten1 (51), we propose that Pot1 stabilizes Stn1-Ten1 recruitment through its interaction with Tpz1 (**Figure 5C**). This idea is supported by the observation that overexpression of *stn1*^*+*^ can restore growth deficiency, telomere loss and telomere elongation of *pot1-1* mutants at restrictive temperature. Upon *pot1* inhibition, Stn1-Ten1 recruitment to telomeres is impaired causing telomere elongation, replication defects and accumulation of ssDNA (**Figure 5C**), all reminiscent of phenotypes observed in *stn1-1* and *stn1-226* mutants (22, 28).

Taken together, our results suggest that Pot1 and Stn1-Ten1 work together not only in controlling the telomerase elongation cycle but also in DNA replication of telomeric and subtelomeric regions, possibly at stalled replication forks. Thus, Pot1, like Stn1-Ten1 and Taz1, is a new component of the Shelterin involved in the DNA replication of telomeres and subtelomeres. Remarkably, the deletion of *rif1* restores viability and telomere loss of *pot1-1* mutant, mimicking the effect observed in *stn1-1* and *stn1-226* mutants. This indicates that the firing of subtelomeric origins can circumvent the replication defects of *pot1-1* by rescuing blocked replication forks.

We established that fission yeast Pot1 is crucial for replication of chromosome ends **(Figure 5C)**. This is achieved by recruiting Stn1-Ten1 and Polα-primase complexes to promote DNA-synthesis upon stalled or blocked replication forks. Unlike mammalian cells, this mechanism appears to be restricted to telomeres since Stn1-Ten function depends uniquely on its interaction with Tpz1/Pot1. The evolutionary difference may have occurred as result of the loss of the CTC1 homolog in fission yeast. In human and mouse cells, POT1 was proposed to promote the telomere lagging strand synthesis by the recruitment of the CST complex (40, 41). Based on our data in fission yeast, we anticipate that human POT1 participates in telomere DNA replication by promoting CST-dependent DNA synthesis at stalled replication forks by preventing the accumulation of RPA-coated ssDNA and repressing ATR activation.

Mammalian CST also fulfils important functions outside of telomeres (52, 54). It facilitates the replication of G-rich regions in the genome (32, 33, 55) and counteracts DNA resection at double strand breaks (DSBs) in cooperation with the Rev7-Shieldin complex (56). The interaction between CST and TPP1-POT1 may not necessary for its function at global genomic G-rich sites or DSBs. However, CST interaction with TPP1-POT1 may be required to promote CST recruitment at the multiple interstitial telomeric sequences (ITS) and facilitate their replication (57). Further research will be necessary to elucidate the role of POT1 in DNA replication of both telomeric and non-telomeric region in mammalian cells.

## Acknowledgments

We thank Dr. Cooper, Dr. Ishikawa, Dr. Bianchi, Dr. McIntosh and Dr. Fleck for providing fission yeast strain. We are grateful to Vincent Géli for reading the manuscript and MGF laboratory for critical comments and discussion. This study was supported by the Portuguese Fundação Ciência e Tecnologia (FCT) project number PTDC/BEX-BCM/5179/14 and the Howard Hughes Medical Institute IECS award to MGF. MGF laboratory is supported by the Université Côte d’Azur - Académie 4 (Installation Grant: Action 2 - 2019) and the Fondation pour la Recherche Médicale (Equipe labélisée FRM, EQU201903007804). PCCB is supported by SFRH/BD/114401/2016 and JME is supported by PTDC/BEX-BCM/5179/14 and PIEF-GA-2013-624759. SC laboratory is supported by the “Ligue Nationale Contre le Cancer” (LNCC) (Equipe labélisée). SC is supported by Project Fondation ARC and by the Agence Nationale de la Research (ANR-16-CE12 TeloMito and ANR-20-CE12 TeloRPA).

## Author contributions

MGF conceived the study and designed the experiments. PCCB performed most experiments. JME performed ChIP experiments. SM performed the DNA 2D-gel experiments. MGF and SC wrote the manuscript, MGF supervised the research.

## Data Availability Statement

The data underlying this article are available in the article and in its online supplementary material.

