## Supplementary material for "Pot1 promotes telomere DNA replication via the Stn1-Ten1 complex in fission yeast": Figure S1

### Supplemental materials

#### Table S1: Fission yeast strains used in this study

| Name | Strain | Genotype | Source |
|---|---|---|---|
| *Wt* | MGF10 | *h- ade6-M210 his3-D1 leu1-32 ura4-D18* | R. McIntosh Lab |
| *pot1-1* | MGF1449 | *h- ade6-M210 his3-D1 leu1-32 ura4-D18 pot1-1-GFP::KanMX6* | J. Cooper Lab |
| *exo1Δ* | MGF293 | *h- ura4-D18 exo1::ura4* | O. Fleck Lab |
| *pot1-1 exo1Δ* | MGF2914 | *h-ade6-M210 his3-D1 leu1-32 ura4-D18 pot1-1-GFP::kanMX6 exo1::hphMX* | This Study |
| *nmt1-3x-stn1* | MGF2377 | *h- ade6-M210 his3-D1 leu1-32 ura4-D18 kanMX6:nmt1-3X-stn1* | MGF10 |
| *nmt1-41x-stn1* | MGF2378 | *h- ade6-M210 his3-D1 leu1-32 ura4-D18 kanMX6:nmt1-41X-stn1* | MGF10 |
| *nmt1-81x-stn1* | MGF2379 | *h- ade6-M210 his3-D1 leu1-32 ura4-D18 kanMX6:nmt1-81X-stn1* | MGF10 |
| *nmt1-3x-stn1 pot1-1* | MGF2949 | *h? ade6-M210 his3-D1 leu1-32 ura4-D18 kanMX6:nmt1-3X-stn1 pot1-1-GFP::kanMX6* | This Study |
| *nmt1-41x-stn1 pot1-1* | MGF2950 | *h? ade6-M210 his3-D1 leu1-32 ura4-D18 kanMX6:nmt1-41X-stn1 pot1-1-GFP::kanMX6* | This study |
| *nmt1-81x-stn1 pot1-1* | MGF2952 | *h? ade6-M210 his3-D1 leu1-32 ura4-D18 kanMX6:nmt1-81X-stn1 pot1-1-GFP::kanMX6* | This study |
| *stn1-1* | MGF2707 | *h- his3-D1 leu1-32 ade6-M210 ura4-D18 stn1-1* | F.Ishikawa Lab |
| *pot1-1 stn1-1* | *MGF2931* | *h+ pot1-1-GFP::KanMX6 stn1-1* | *This study* |
| *pot1-1 rif1Δ* | *MGF2924* | *h+ pot1-1-GFP::KanMX6 rif::hph* | *This study* |
| *stn1-1 rif1Δ* | MGF2954 | *h- ade6-M21? his3-D1? leu1-32 ura4-D18 stn1-1 rif1::hphMX6* | This study |
| *Wt + pREP41-empty* | MGF2665 | *h? ade6-M210 his3-D1 leu1-32 ura4-D18 pREP41-nmt41-empty-Leu2* | MGF10 |
| *Wt + pREP41-pol1* | MGF2666 | *h? ade6-M210 his3-D1 leu1-32 ura4-D18 pREP41-nmt41-pol1-Leu2* | MGF10 |
| *pot1-1 + pREP41-empty* | | *h+ ade6-M210 his3-D1 leu1-32 ura4-D18 pot1-1-GFP::kanMX6 pREP41-nmt41-empty-Leu2* | This Study |
| *pot1-1 + pREP41-pol1* | | *h+ ade6-M210 his3-D1 leu1-32 ura4-D18 pot1-1-GFP::kanMX6 pREP41-nmt41-pol1-Leu2* | This Study |

### Figure S1

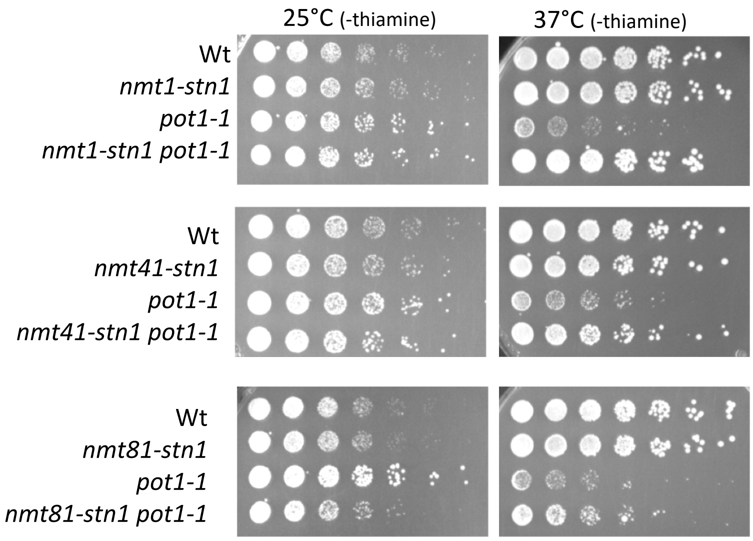


**Figure S1. A.) Overexpression of Stn1 rescues pot1-1 cells viability.** Different levels of Stn1 overexpression in *pot1-1* cells were tested and all of them rescued *pot1-1* cells viability. Spot Assay was performed in PMG solid media which lacks thiamine and incubated at 25°C to 37°C.
